# Spectrum of protein localization in proteomes captures evolutionary relation between species

**DOI:** 10.1101/845362

**Authors:** Valérie Marot-Lassauzaie, Tatyana Goldberg, Burkhard Rost

## Abstract

The native subcellular localization or cellular compartment of a protein is the one in which it acts most often; it is one aspect of protein function. Do ten eukaryotic model organisms differ in their *location spectrum*, i.e. the fraction of its proteome in each of its seven major compartments? As experimental annotations of locations remain biased and incomplete, we need prediction methods to answer this question. To gauge the bias of prediction methods, we merged all available experimental annotations for the human proteome. In doing so, we found important values in both Swiss-Prot and the Human Protein Atlas (HPA). After systematic bias corrections, the complete but faulty prediction methods appeared to be more appropriate to compare location spectra between species than the incomplete more accurate experimental data. This work compared the location spectra for ten eukaryotes: *Homo sapiens, Gorilla gorilla, Pan troglodytes, Mus musculus, Rattus norvegicus, Drosophila melanogaster, Anopheles gambiae, Caenorhabitis elegans, Saccharomyces cerevisiae* and *Schizosaccharomyces pombe*. Overall, the predicted location spectra were similar. However, the detailed differences were significant enough to plot trees and 2D (PCA) maps relating the ten organisms using a simple Euclidean distance in seven states, corresponding to the seven studied localization classes. The relations based on the simple predicted location spectra captured aspects of cross-species comparisons usually revealed only by much more detailed evolutionary comparisons.

## Introduction

### Location spectrum of an organism

Eukaryotic cells contain many distinct compartments separated by membranes. This separation allows to create functionally specialized spaces with slightly different biophysical features [1]. The atlas of where proteins predominantly perform their function, i.e. their native localization or compartment therefore contains important information about protein function that is used to classify function in the Gene Ontology (GO) [2]. We refer to the *<u>location spectrum</u>* as to the percentage of proteins in each location for an entire organism; one component of this spectrum is e.g. the fraction of secreted proteins. Does anything as simple and abstract as the location spectrum contain any relevant information about an organism?

Previous experimental studies have attempted to determine the proteome-wide location spectrum for model organisms such as baker’s yeast (*Saccharomyces cerevisiae*) [3] and *Pseudomonas aeruginosa* [4]. Despite more than a decade of such successful studies, only 75% of yeast are classified into one of 22 locations. Furthermore, we show here that the location spectrum for human cannot be estimated using the experimental-only annotations (Fig. SOM3, Supporting Online Material cf. Results).

### Most proteins have one dominant “native” localization

Many proteins “travel”, i.e. they function in more than one localization over the course of their existence. Most proteins, however, have one pre-dominant, native localization to accomplish their function as suggested by the following argument. Assume the opposite, namely that each protein is equally often observed in D different localizations. If we forced annotations to be limited to a single localization (i.e. ignore all annotations we find other than the first) and we compared two sets of annotations of localization (from predictions or experiments) that are essentially error-free, comparing 1000 proteins, we would find 1000/D to agree between the methods due to picking only one of D. Some analysis of data for human might be misunderstood to suggest D=3 (Results). For this number 1000/3=333 proteins would agree due to the combination of error-free annotations and picking only the first annotation. Assuming L classes of localization (for simplicity let L be 10), then one tenth of the 667 disagreeing (1000-333) proteins would match at random, i.e. another 68, bringing the total to 401. In other words, the agreement between two error-free data sets would appear to be 40%. For D=2 (two native localizations per protein), this number would rise to 55%, for D=1.5 to 78%. However, good prediction methods reach levels of performance above 65% in 18 states, and above 80% for the best classes [5] even when using an annotation data set that assumes only one localization to be correct. Thus, this little Gedankenexperiment refutes the hypothesis that most proteins have more than one “native” localization.

### Annotating localization in human proteome

A more complex organism with many readily available experimental location annotations is *Homo sapiens*. Assuming a predominant native localization, we analyzed the agreement between experimental results, then combined results into a large set with reliable annotations, and analyzed how much the given experimental annotations reveal about the expected location spectrum in an organism. We found that inaccurate predictions that are available for all proteins provided a much better proxy for the real location spectrum of an organism than much more accurate but incomplete experimental observations. In order to really establish this result, we needed to correct the bias introduced by prediction methods that predict some locations better than others. Given this error correction, we predicted the localization spectrum for ten model organisms. Surprisingly, the comparison of the spectrum of locations alone was enough to reproduce the evolutionary similarities between the organisms. This finding indicates that this measure contains relevant information specific to each organism.

## Methods & Materials

### Proteomes

The sequences for the reference proteomes were taken from the EMBL-EBI database (https://www.ebi.ac.uk/reference_proteomes) [6]. The Human proteome was taken from the 4^th^ release of 2016 which contains 21,018 proteins. For the cross-species comparison nine reference organisms were chosen from release 2017-4.

### Experimental annotations

Experimental annotations for the localization of human proteins were taken from Swiss-Prot [7] and The Human Protein Atlas (HPA) [8]. This resulted in a set of 5,563 proteins with experimental annotations from Swiss-Prot (release 2017_1; human proteome release up000005640), and in 12,036 from The Human Protein Atlas (version 15; confined to 32 localization classes). We had access to one additional set of experimental annotation in GO format extracted from scientific literature by the tool LocText [9].

### Prediction methods

The following five prediction methods were compared for human. <u>LocTree2</u> [5] uses profile kernels Support Vector Machines (SVMs) through a decision tree; it predicts one of 18 localizations for eukaryotes. <u>LocTree3</u> [10] combines LocTree2 with homology-based inference (in as many different classes as are experimentally annotated). <u>Hum-mPloc3.0</u> [11] predicts 12 localization classes by combining residue-based statistical features, with conserved domains and Gene Ontology annotations. Several classes are predicted for each protein; only the one with the highest score was kept. <u>MultiLoc2</u> [12] predicts 11 localization classes by integrating the output of six sequence based classifiers (SVMTarget, SVMSA, SVMaac, MotifSearch, GOLoc, PhyloLoc) through a final SVM. <u>WoLF PSort</u> [13] predicts 10 localization classes by first converting a sequence into features indicative of localization (amino acid composition, sorting signals and functional motifs). A k-nearest neighbor classifier is applied to those features to predict.

The five prediction methods were evaluated against two reference sets of proteins with known annotations that were most likely not sequence similar to any protein that had been used to develop and asses the prediction methods applied. The first reference set was taken from the HPA (published after the development of the methods). The second reference set was extracted from scientific publications using LocText [9]. None of those had been annotated in Swiss-Prot. To avoid further complications, only proteins with a single annotation were kept. This resulted in a set of 2,000 proteins with reliable experimental annotations from HPA and 1,315 with less reliable maps to annotations from LocText (less reliable due to possible mistakes in the text mining).

### Comparing predictions

Since the prediction methods predict a different number of classes, those predictions needed to be made comparable by reduction to seven main localization classes shared between all tools (SOM: Supporting Online Materials Table S1 for mapping): secreted, nucleus, cytoplasm, plasma membrane, mitochondrion, endoplasmic reticulum (ER) and Golgi apparatus. All proteins for which the experimental annotations could not be mapped to those seven classes were excluded from the analysis.

### Error corrections for estimates of distributions

Prediction methods tend to make specific mistakes, and these mistakes are not the same for all classes. Therefore, the predicted spectrum of locations for an entire organism cannot be estimated accurately enough from using the output of prediction methods directly. Instead, the specific errors have to be corrected [14]. Toward this end, the confusion matrices for each prediction method needed for the correction were built by establishing the performance for human proteins with a single experimental annotation in Swiss-Prot.

### Distance between distributions

For human proteins, the 7-state distributions directly obtained from the experimental annotations were compared with the ones predicted by the five selected prediction tools. The straightforward Euclidean distance for this 7-state distribution served as proxy for the difference between predictions and experiments. The distance d between two points *p=(p*_*1*_,*p*_*2*_,…,*p*_*n*_) and *q=(q*_*1*_,*q*_*2*_,…,*q*_*n*_) in a n-dimensional state is given by the Pythagorean formula:

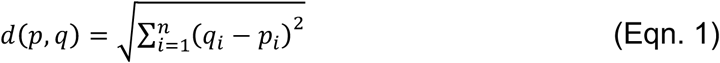

Here, the seven dimensions were given by the location spectrum in seven main subcellular localization classes. For the other nine model organisms, the same Euclidean distance established how similar those organisms were according to the predicted spectrum of locations. These distances were visualized in two different ways. Firstly through a UPGMA [15] tree built using the R-package phangorn [16]. Secondly, through a 2D view originating from the multidimensional scaling representation of the distances with the stats R-package [17].

### Error-correction using confusion matrix

As described previously [14], we corrected the predicted location spectra through the performance estimates given in the confusion matrix M_p,o_ (TP true positives, TN true negatives, FP false positives and FN false negatives), where the elements Mp,o give the number of proteins predicted in state p and observed in state o. The diagonal contains the correct predictions with p=o. From M we can compute a new n*n matrix M’ with:

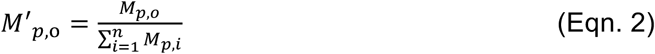

This new matrix provided the ratio by which each location was normalized. The predictions for an entire data set P=(p_1_,p_2_,…,p_n_) were corrected to P_c_=(p_1_,p_2_,…,p_n_) with 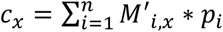.

## Results and Discussion

### Reliable experimental location annotations for 37% of all human proteins

Both Swiss-Prot [7] and The Human Protein Atlas (HPA) [8] experimentally annotate sub-cellular localization of proteins by many different criteria. HPA exclusively uses experimental annotations of varying quality (Validated > Supportive > Uncertain > Unreliable). The evidence codes added to Swiss-Prot clearly distinguish experimental annotations from those inferred through sequence comparisons. We only considered annotations with explicit experimental evidence (evidence code ECO:0000269). If both databases were error-free, their annotations would agree independently of the experimental sources. For this analysis, we only considered the subset of the proteins for which both Swiss-Prot and HPA had experimental annotations. Annotations were counted as identical when any location matched, e.g. a protein with the Swiss-Prot annotation “nucleus, cytoplasm” and the HPA annotation “cytoplasm, mitochondria, Golgi” was considered to have identical annotations. In this way, 94% of the HPA annotations with higher reliability (HPA-level: *Validated* and *Supportive*: 2261 cases, Table 1) agreed with Swiss-Prot, while only 54% of those with lower reliability did (HPA-level: *Uncertain* and *Unreliable*: 909 cases; Table 1; Fig. 1). We considered the first value (94%) high enough to label those annotations as “*reliable*”, all the others as “*speculative”* (Table 1). Overall, HPA and Swiss-Prot together have reliable experimental information for 7,705 of the human proteins (37% Fig. 1: 2572+2261+1963+909), and both share their reliable annotations for only 2,261 (11%) proteins (at least one annotation in common between Swiss-Prot and HPA *reliable*). Thus, a simple merge of the two databases added considerably to each of these outstanding resources.

**Table 1:** Reliable Human Protein Atlas (HPA) annotations agree with Swiss-Prot*.

| *HPA level* | *Nprot* | *Any HPA annotation identical to any in Swiss-Prot* | *Expected at random* |
| --- | --- | --- | --- |
| *Validated* | 644 | 95% | 38% |
| *Supportive* | 1617 | 94% | 39% |
| *Uncertain* | 783 | 57% | 45% |
| *Unreliable* | 126 | 39% | 44% |
| *Merged used here* |  |  |  |
| *Reliable* | 2261 | 94% | 39% |
| *Speculative* | 909 | 54% | 45% |

**Figure 1:**
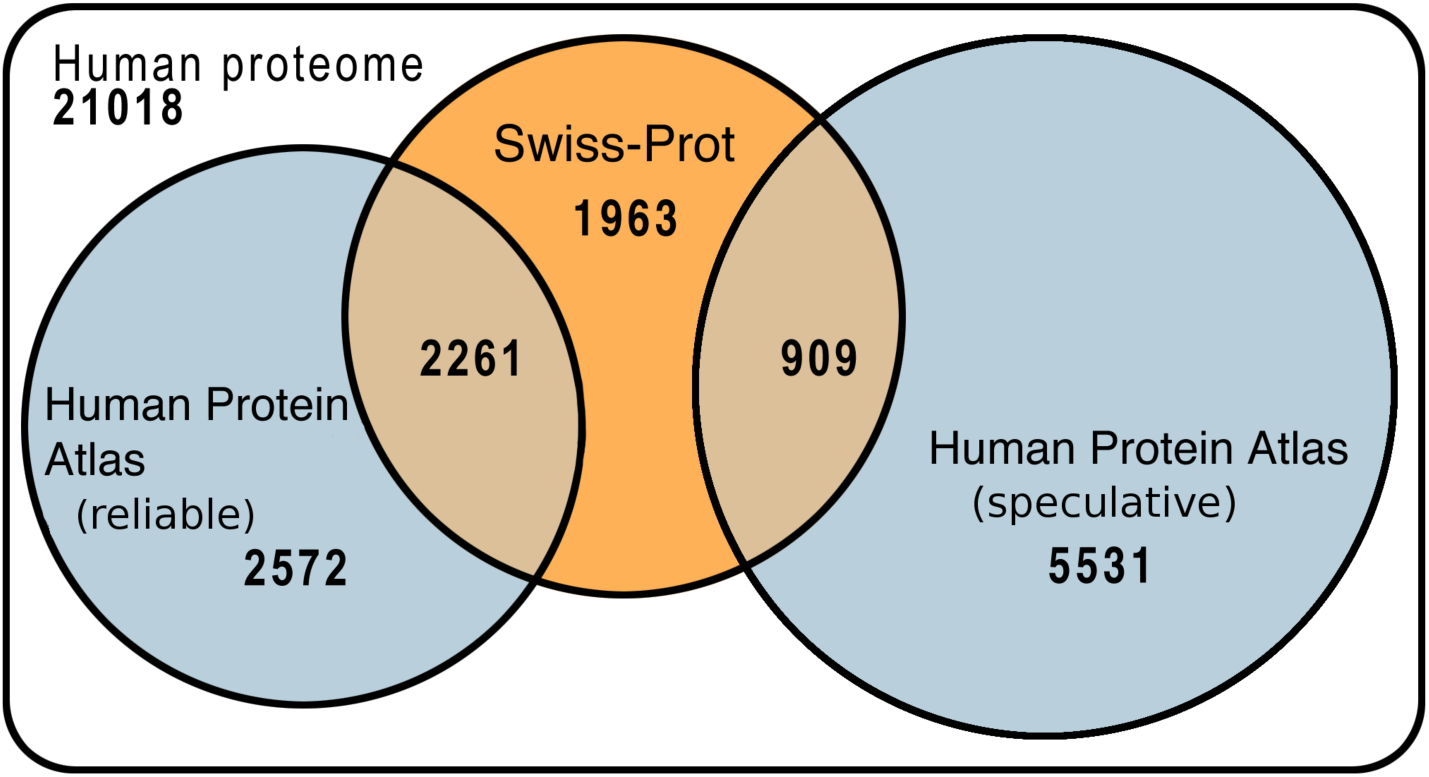
Protein localization in human proteome annotated by experiments. The Venn-diagram compares experimental annotations of human proteins between Swiss-Prot [7] and The Human Protein Atlas (HPA) [8]. For ease of visibility the size of the white background (entire human proteome with 21,018 proteins) is not to scale. HPA provides four levels in the reliability of their annotations. We grouped those into *reliable* (HPA keywords: *validated* and *supportive*, 94% agreement with Swiss-Prot: Table 1) and *speculative* (HPA keywords: *uncertain* and *unreliable*, 54% agreement with Swiss-Prot: Table 1). The agreement between HPA *speculative* and Swiss-Prot is only slightly above random (Table 1). For instance, 2,261 proteins have HPA have reliable annotations and match at least one of the experimental Swiss-Prot annotations (evidence code: ECO:0000269) while 37% of all human proteins (7705=1963+2572+2261+909) have reliable experimental annotations (either Swiss-Prot ECO:0000269 or HPA validated and supportive).

### HPA also provides many speculative annotations

While the HPA top level *validated* agreed to 95% with Swiss-Prot and the HPA level *supportive* to 94%, the agreement dropped below 60% for the HPA level *uncertain* (57%) and to 39% for *unreliable* (Table 1). Considering over 90% agreement sufficient to consider the annotation as “reliable”, we grouped the top two HPA levels into a class “reliable” (as opposed to “speculative” for the lower two, Table 1) and used only reliable annotations to assess methods. These findings underline that the authors of HPA have done a very good job at correctly distinguishing between more and less reliable results.

* *<u>HPA level:</u>* levels of reliability provided by The Human Protein Atlas (HPA) [8]; *<u>Nprot:</u>* number of human proteins included in comparison (note: restricted to proteins from HPA with a match in Swiss-Prot); *<u>Merged used here</u>*: according to the agreement between HPA and Swiss-Prot, the two best HPA levels were grouped into “*reliable*”, the two worst into “*speculative*”; *<u>Any HPA annotation identical to any in Swiss-Prot:</u>* percentage of proteins for which Swiss-Prot (experimental only [7]) annotations agree with at least one of the localization classes of HPA (e.g. HPA “cytoplasm, nucleus” was considered *identical* to Swiss-Prot “cytoplasm, plasma membrane”); *<u>Expected at random:</u>* agreement between annotations when randomly shuffling annotations (in more detail: randomly pick proteins from the Swiss-Prot set, compare to the HPA proteins of the corresponding HPA level, repeat 100 times for each level and take the mean).

The HPA *speculative* class still matched Swiss-Prot at levels higher than expected by chance (Table 1: 54% vs. 42%; note the original HPA level *unreliable* appeared less similar to Swiss-Prot than random). However, this match score was clearly lower than that for prediction methods.

### Localization reliably inferred for 81% of the human protein families

Proteins with similar sequences tend to have similar function, hence also similar localization [18]. For instance, >92% of the sequence similar pairs of proteins have the same localization [18] at an HVAL>4 (corresponding to 24% percentage pairwise sequence identity for alignments spanning ≥250 residues [19]). We applied this threshold to cluster all human proteins through UniqueProt [20]. This resulted in 3,148 *families* (defined as HVAL>4 for all proteins in that *family* to a representative seed protein). 1,920 of those (62%) were covered by experimental annotation from at least one protein within the family. Thus, homology-based inference enables the prediction of localization for all family members at over 90% accuracy. However, those families tended to be larger than those that were not covered. Therefore, homology-based inference covered 18,840 (89%) of all human proteins.

Despite this high coverage, homology-based inference alone could not estimate the full “spectrum of localizations” in an organism. If we used the combined annotations of both databases to infer the locations of the 1,920 families, the number of multiple locations would be too high: 16,378 of the 18,840 proteins were predicted by homology in multiple compartments (on average 3.3 compartments per protein). If true, our comparisons between Swiss-Prot and HPA could never have reached above 40% agreement (argument in Introduction: 1/3=33% + random=40%). Not even half the value of what the top HPA annotations actually reached (class: *reliable*=94% Table 1). Another non-sense consequence is the following. If using only experimental and homology-inferred annotations, we would predict 15,169 human proteins (72%) to be nuclear. Whatever the true value might be, clearly 72% is much too large. Although the corresponding number for families is slightly lower (68% SOM: Fig. S1), it still appears unlikely. In terms of per-family view even more misleading is the number of 49% of all human protein families associated with the plasma membrane (Fig. S1) when fewer than 25% of all human proteins have a membrane helix, and only 13% have more than one transmembrane helix [21]. Thus, we apparently cannot estimate the spectrum of localizations for an organism from experimental and homology-inferred annotations alone.

Another limitation of using experimental annotations enriched by homology-based inference alone to estimate the spectrum of locations in human was highlighted by monitoring the number of families in multiple locations: 16,378 of the 18,840 proteins reachable by “experiment + homology” were found in multiple compartments (on average 3.3 compartments per protein). Following the argument made in the Introduction (D=3.3), this would imply a maximal performance of a 5-state prediction method of 44%, i.e. much lower than observed.

### Accurate predictions of location spectrum for organisms

Overall, most prediction methods came relatively close (Fig. 2A: the more white the fewer the mistakes) to estimating the 7-class *location spectrum* for the new data sets that had not been used for developing the methods (HPA: new data from The Human Protein Atlas, and the *LocText* text mining results [9]). When error-correcting the predicted spectra according to the performance confusion matrix [14], the observations and predictions became more similar for most methods (Fig. 2B; exceptions LocTree3 and WolfPsort for HPA;). After error correction Hum-mPloc3.0 estimated the location spectrum of the experimental data best (Fig. 2B: most white for HPA column on left). However, since this tool is limited to human, we had to continue our cross-organism analysis with the second-best prediction, namely the error-corrected version of *LocTree3*.

**Figure 2:**
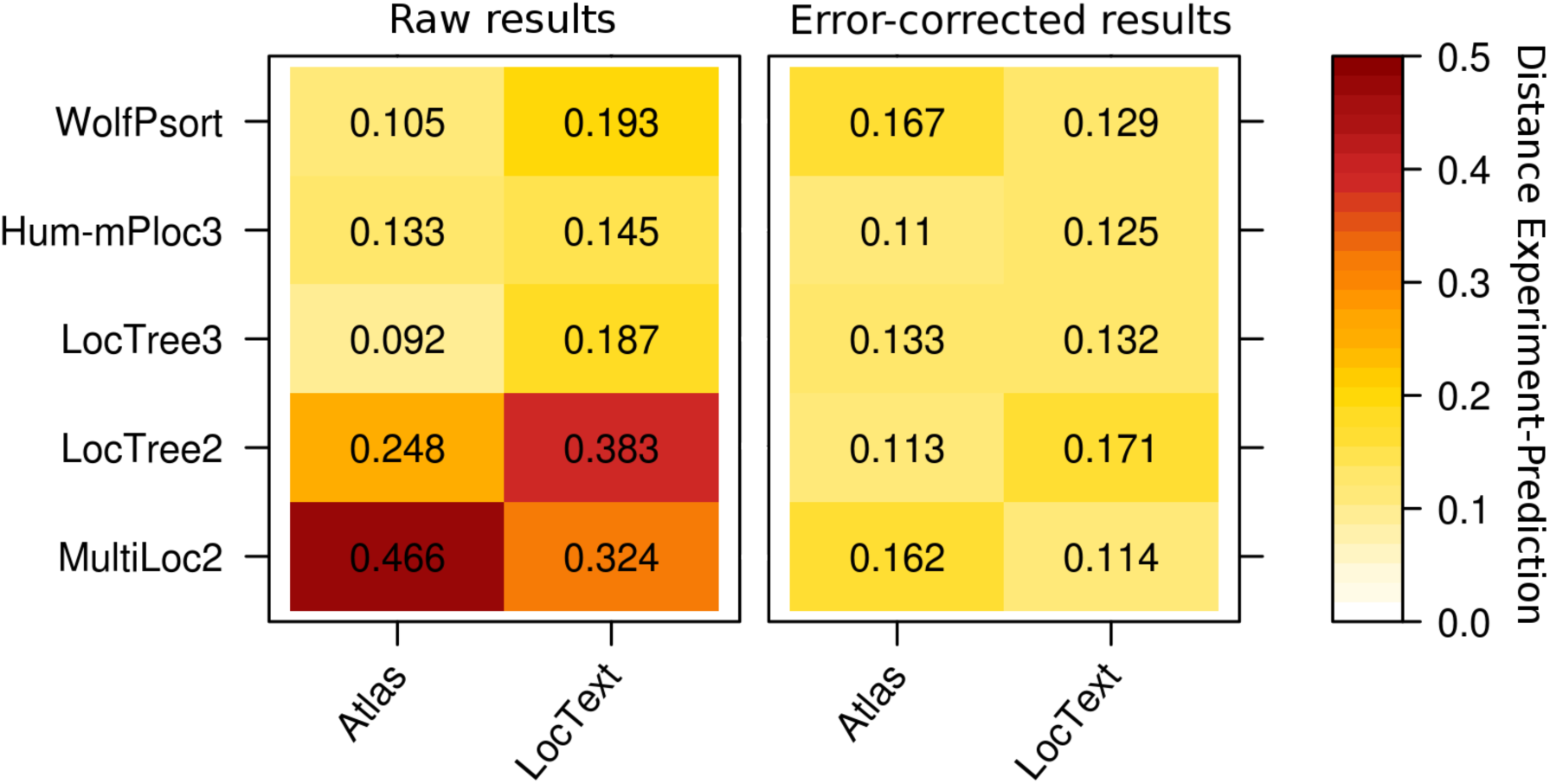
Agreement in location spectrum for prediction methods. For two different sets of proteins that were not used by the prediction methods (HPA and LocText), the heat maps give the Euclidian distances of the location spectra (Eqn. 1) for several prediction methods. The left panel (*Raw results*) show the results directly obtained from the prediction methods, the right panel (*Error-corrected results*) show the same after the application of a simple error-correction using the confusion matrix for each method (Eqn. 2 and 3) [14]. A darker, redder color indicates a higher distance, i.e. a worse prediction. Almost all methods improve through the error corrections and almost all estimate the location spectra very accurately.

### Location spectra capture aspects in the evolution of organisms

Predictions of the location spectra for ten completely sequenced model eukaryotes in seven main localization classes were computed by *LocTree3* (Fig. 3, SOM: Table S2). The predicted location spectra were largely similar between the ten proteomes. However, the small but significant differences sufficed to draw UPGMA-trees relating those organisms that appeared reasonable in the following key aspects (Fig. 3.A.). (1) The two yeast types were grouped together and separated from the multicellular organisms. (2) Mammals were grouped together. (3) The two rodents were separated from the three apes. (4) Furthermore, *Pan troglodytes* and *Homo sapiens* were considered to be closer to each other than to *Gorilla gorilla*.

**Figure 3:**
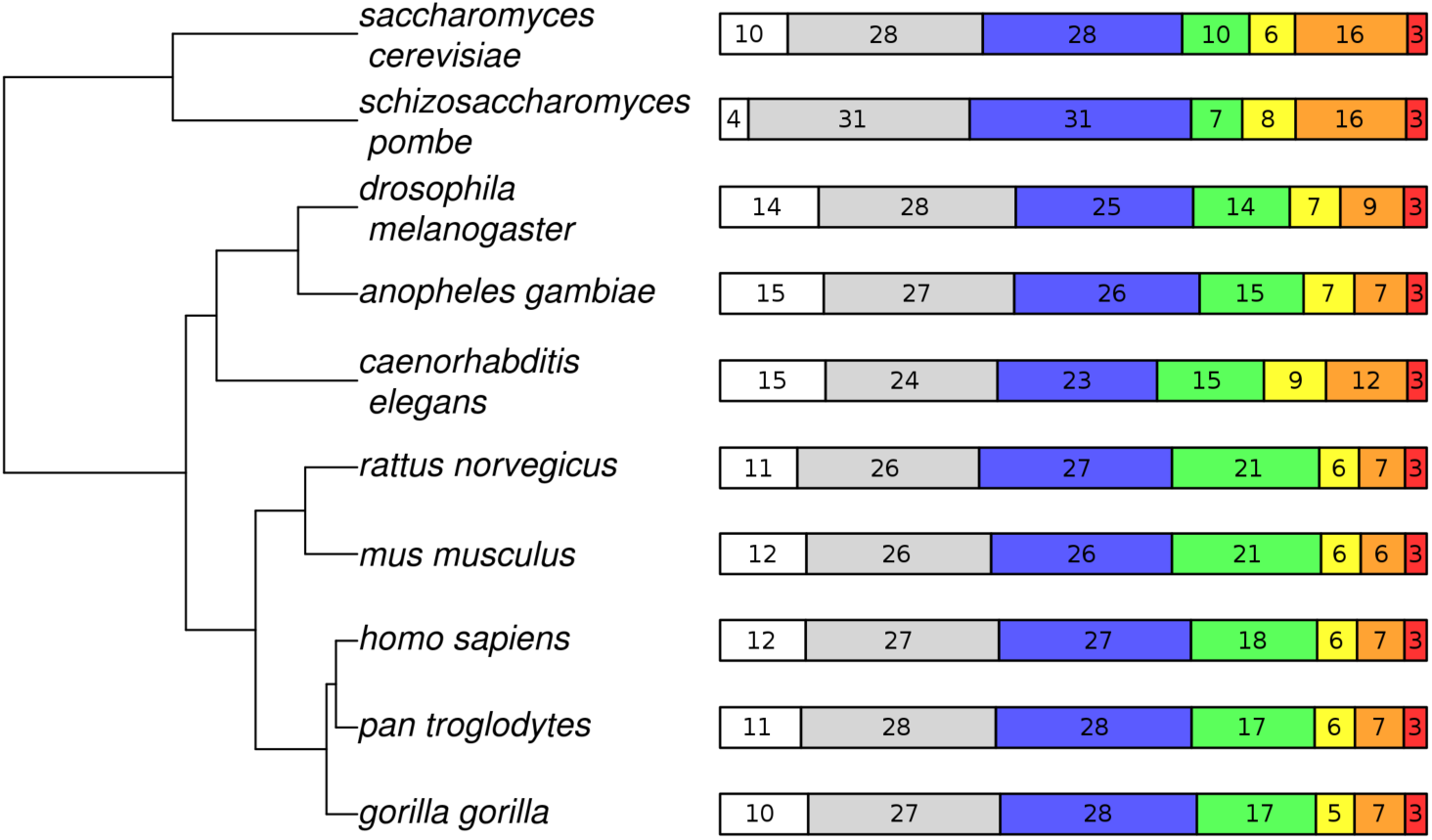
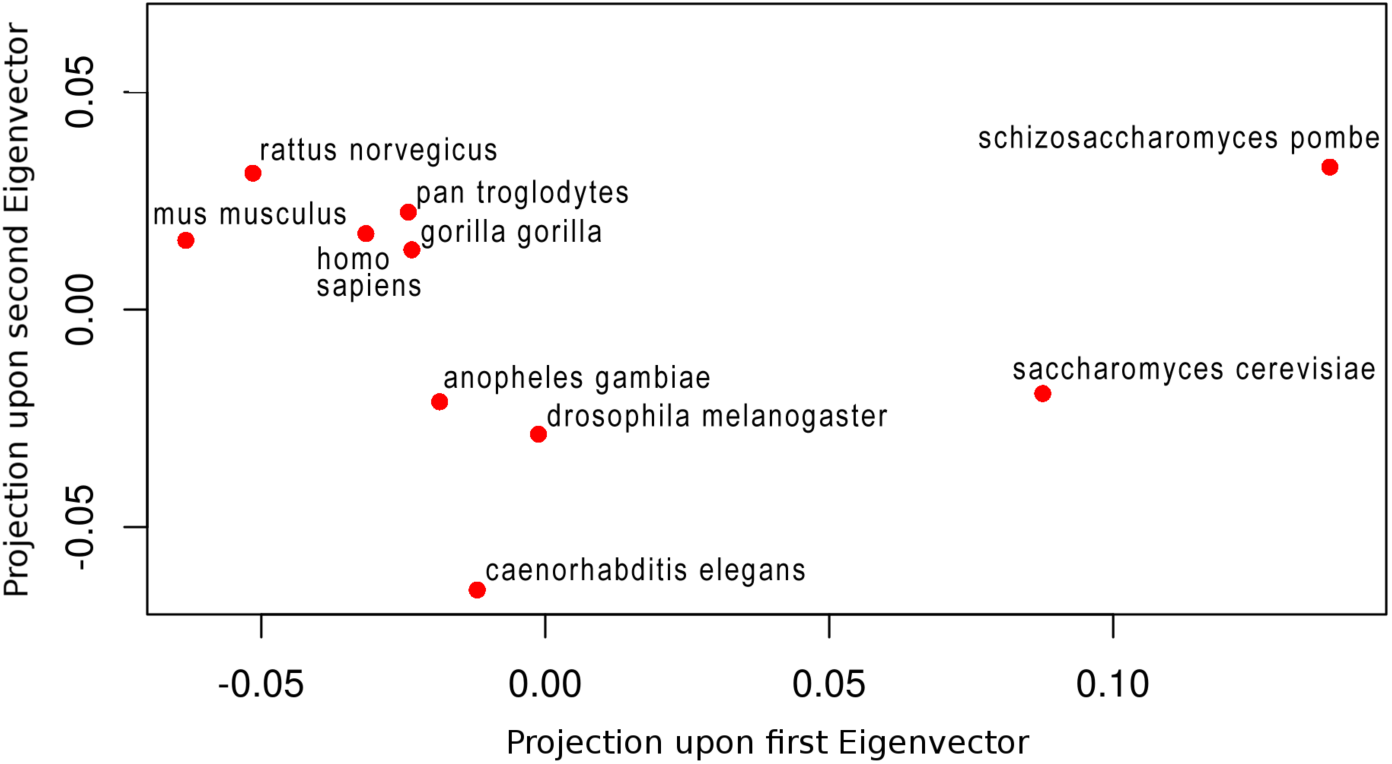
Grouping of ten eukaryotes according to predicted *location spectra*. We computed the Euclidean distances (Eqn. 1) between the proteome-wide distributions predicted by *LocTree3* with error-correction (Eqn. 2 and 3) [14] for each of the ten reference organisms. Those values were plotted onto a UPGMA tree (top panel A) and shown through PCA in 2D (lower panel B). **(A)** UPGMA tree along with a bar representing the predicted distribution in the seven main subcellular localization classes is shown for each organism. The seven localization classes (from left to right): secreted (white), nuclear (gray), cytoplasmic (blue), plasma membrane (green), mitochondrial (yellow), endoplasmic reticulum (orange) and Golgi apparatus (red). Despite the small differences, the resulting tree largely agrees with what we would expect from evolution. **(B)** The PCA adds more details to the comparison between species. Two interesting aspects are the large differences between the two yeast-species (y-axis) and the approximate triangle between mouse, rat and human.

The tree representation simplifies and thereby might miss important relations. Therefore, we also displayed the relations in two dimensions (2D) through a simple principle component analysis (PCA; first two eigenvector projections shown in Fig. 3.B.). The 2D view confirmed the principle findings from the tree and added interesting details. For instance, in 2D the two yeast types remain separated from all multi-cellular organisms, but are much more separated from each other on the y-axis (2^nd^ eigenvector) than for instance the apes from each other. This could be due to the smaller size of their genome comparatively to the other organisms or to the short generation time which allows for a faster divergence. Another interesting observation was that the proximity on the y-axis (2^nd^ eigenvector) between both rodents (mouse and rat) was similar to the proximity of each rodent to human. Here again, this observation could be explained by the shorter generation time of these organisms allowing for a comparatively high distance.

## Conclusions

We showed that experiments enriched by homology-based inference accurately annotate the subcellular localization for 81% of all human proteins (Fig. 1). Nevertheless, prediction methods are much better at estimating the location spectrum for entire proteomes, i.e. the composition of proteins in each of the major compartments. In fact, through the application of a simple method for correcting prediction-mistakes, the predicted spectra appeared to be very accurate (Fig. 2). Applying such an error-corrected whole proteome prediction with LocTree3 to ten model organisms suggested rather similar location spectra (Fig. 3). Nevertheless, the difference even allowed to build a tree based on these spectra that reflected aspects of the evolution between those organisms both on the level of a 1D tree (Fig. 3) and of a 2D proximity map (Fig. 4). Overall, this clearly appeared to confirm the validity of the approach to estimating the spectrum of location through prediction methods.

## Abbreviations used

3D: three-dimensional;
3D structure: three-dimensional coordinates of protein structure;
PCA: principal component analysis;
HPA: human protein atlas;
ER: endoplasmic reticulum

## Acknowledgements

Thanks to Tim Karl (TUM) for invaluable help with hardware and software; to Inga Weise (TUM) for administrative support; to Michael Bernhofer & Jonas Reeb (both TUM) for help with the error correction; to Maria Schelling and Michael Bernhofer for advice and help with the validation. Thanks to Ioannis Xenarios (Swiss-Prot, SIB, Geneva), Mathias Uhlen (HPA Uppsala Univ), and their crews for maintaining excellent databases and to all experimentalists who enabled this analysis by making their data publicly available.

