## Supplementary materials: table S1, table S2 and figure S1 for "Spectrum of protein localization in proteomes captures evolutionary relation between species"

### Supporting online material (SOM) for: Correcting mistakes in predicting protein-wide distributions

**Valérie Marot-Lassauzaie, Michael Bernhofer & Burkhard Rost**

#### Short description of Supporting Online Material

This file contains the supporting online material (SOM) for the publication "Correcting against mistakes in predicting protein-wide distributions".

• **Table S1** shows how the output from the three prediction tools and from The Human Protein Atlas [1] was converted to a set of seven main localization classes for the comparison of the output.

• **Table S2** gives the error-corrected number of proteins predicted in each of the seven localization classes in each of the ten model organisms.

• **Fig. S1** demonstrates through a racetrack plot how the prediction of localization for the human proteome through homology based inference would generate a rather misleading localization spectrum.

### Material

#### Table S1: Conversion of predictions and HPA to seven localization classes ◊.

| *Localization class* | *The Human Protein Atlas (HPA) [1]* | *MultiLoc2 [2]* | *Hum-mPloc3.0 [3]* | *LocTree2 [4]* |
|---|---|---|---|---|
| Secreted | N/A | Extracellular | Extracellular | Extracellular |
| Nucleus | nucleoplasm, nuclear bodies, nuclear speckles, nucleus, nucleoli, nucleoli fibrillar centre, nuclear membrane | Nucleus | Nucleus | Nucleus, nucleus membrane |
| Cytoplasm | Cytosol, microtubule, microtubule ends, microtubule organising centre, intermediate filaments, cytoplasmic bodies, actin filaments, centrosome, midbody, cytokinetic bridge, mitotic spindle, rods & rinds, midbody ring | Cytoskeleton, Cytoplasmic | Centriole, Cytoplasm, Cytoskeleton | Cytosol |
| Plasma membrane | Plasma membrane, Focal adhesion sites, cell junction | Plasma membrane | Plasma membrane | Plasma membrane |
| Mitochondrion | Mitochondrion | Mitochondrion | Mitochondrion | Mitochondria, mitochondria membrane, chloroplast[1], chloroplast membrane[1], plastid[1] |
| Endoplasmic reticulum | Endoplasmic reticulum | Endoplasmic reticulum | Endoplasmic reticulum | Endoplasmic reticulum, endoplasmic reticulum membrane |
| Golgi apparatus | Golgi apparatus | Golgi apparatus | Golgi apparatus | Golgi apparatus, Golgi apparatus membrane |

◊ Conversion of the output from the prediction tools (MultiLoc2 [2], LocTree2 [4], Hum-mPloc3.0 [3]) and from The Human Protein Atlas [1] to the seven main localization classes used for the comparison of the outputs. For each tool or resource, the mapping from the tool's output classes to the corresponding reference class is shown. Any entry not fitting in the seven reference classes was excluded from comparison.

1: for LocTree2 [4] “Chloroplast and plastid classes are valid for plant proteins only, therefore, if the origin of a non-plant protein is known, please consider prediction of these classes as mitochondrial”

---

**Table S2: Location spectra predicted for ten eukaryotes proteomes ◊.**

| Organism | Secreted | Nucleus | Cytoplasm | Plasma membrane | Mitochondrion | Endoplasmic reticulum | Golgi apparatus |
|---|---|---|---|---|---|---|---|
| HUMAN | 2520 | 5686 | 5648 | 3710 | 1174 | 1395 | 656 |
| ANOGA | 1701 | 3128 | 3044 | 1707 | 834 | 864 | 324 |
| CAELL | 2958 | 4818 | 4479 | 2995 | 1714 | 2282 | 542 |
| DROME | 1876 | 3750 | 3377 | 1839 | 962 | 1208 | 435 |
| GORGO | 2585 | 5637 | 5772 | 3495 | 1135 | 1483 | 636 |
| MOUSE | 2693 | 5759 | 5637 | 4651 | 1239 | 1389 | 667 |
| PANTR | 2153 | 5177 | 5201 | 3272 | 1069 | 1300 | 609 |
| RAT | 2315 | 5452 | 5783 | 4417 | 1192 | 1378 | 653 |
| SCHPO | 201 | 1569 | 1571 | 363 | 378 | 784 | 146 |
| YEAST | 625 | 1802 | 1842 | 621 | 422 | 1038 | 178 |

◊ Number of proteins predicted in each of the seven sub-cellular localisations (after error correction [5]) for the ten model organisms: *Homo sapiens* (HUMAN), *Drosophila melanogaster* (DROME), *Anopheles gambiae* (ANOGA), *Rattus norvegicus* (RAT), *Mus musculus* (MOUSE), *Pan troglodytes* (PANTR), *Gorilla gorilla* (GORGO), *Caenorhabditis elegans* (CAEEL), *Saccharomyces cerevisiae* (YEAST) and *Schizosaccharomyces pombe* (SCHPO). Predictions were computed by LocTree3.

**Fig. S1**

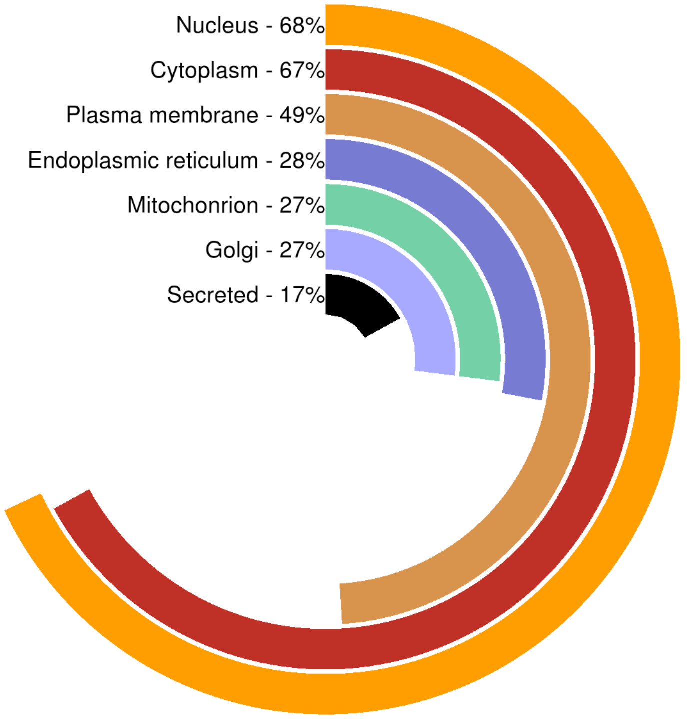


**Fig. S1: Racetrack plot showing the prediction of localization for the human proteome through homology based inference.** Using UniqueProt [6], we clustered all human proteins at threshold HVAL>4. This resulted in 3,148 families. 1,920 families were covered by experimental annotation in HPA or Swiss-Prot [7] from at least one protein in the family. Considering all proteins in a family to have the same localization as the annotated proteins of the family, we could infer the localization of 18,840 (89%) human proteins. The racetrack plot clearly illustrates limits of such an approach. For instance, it is very unlikely that 68% of all families of human proteins are nuclear, and equally unlikely that the roughly 25% of the human transmembrane proteins constitute 49% of all families, in particular given that most membrane families are unusually large.
